# Multispecies coalescent analysis unravels the non-monophyly and controversial relationships of Hexapoda

**DOI:** 10.1101/187997

**Authors:** Lucas A. Freitas, Beatriz Mello, Carlos G. Schrago

## Abstract

With the increase in the availability of genomic data, sequences from different loci are usually concatenated in a supermatrix for phylogenetic inference. However, as an alternative to the supermatrix approach, several implementations of the multispecies coalescent (MSC) have been increasingly used in phylogenomic analyses due to their advantages in accommodating gene tree topological heterogeneity by taking account population-level processes. Moreover, the development of faster algorithms under the MSC is enabling the analysis of thousands of loci/taxa. Here, we explored the MSC approach for a phylogenomic dataset of Insecta. Even with the challenges posed by insects, due to large effective population sizes coupled with short deep internal branches, our MSC analysis could recover several orders and evolutionary relationships in agreement with current insect systematics. However, some phylogenetic relationships were not recovered by MSC methods. Most noticeable, a remiped crustacean was positioned within the Insecta. Additionally, the interordinal relationships within Polyneoptera and Neuropteroidea contradicted recent works, by suggesting the non-monophyly of Neuroptera. We notice, however, that these phylogenetic arrangements were also poorly supported by previous analyses and that they were sensitive to gene sampling.

## Introduction

Biological sciences have been experiencing a boom in the availability of genomic data, so that the number of loci employed to reconstruct phylogenies has increased significantly in the last decade. When dealing with multiple genomic regions, the most usual approach employed by authors consists in the concatenation of such aligned sequences into a supermatrix to build the phylogeny.

Thus far, the main microevolutionary phenomenon accounted for at species tree estimation methods is incomplete lineage sorting (ILS), which is mathematically treated as modeled by the multispecies coalescent (MSC) (Degnan and Rosenberg, 2009). MSC is a derivation of the coalescent theory (Kingman, 1982) that allows the coalescent process to occur independently along branches of the phylogeny. It models gene tree topology heterogeneity, as different *loci* are treated as independent trials of the coalescent processes within the phylogeny. The fact of taking into account the discordant information derived from a sample of gene trees is the principal advantage of MSC over supermatrix methods, which assume the same history for all genes (Edwards, 2009).

Recent studies applied the MSC to infer the relationships of several animal groups (Cannon et al., 2016; Jarvis et al., 2014; Lanier and Knowles, 2015; Song et al., 2012; Xi et al., 2014). However, due to the computational limitation of MSC algorithms, the datasets employed were limited to dozens of species and few hundreds of loci (Zimmermann et al., 2014). In this sense, the advent of faster MSC methods, such as ASTRAL (Mirarab et al., 2014), enables MSC-based phylogeny estimation using thousands of species/loci. Importantly, it allows the use of genomic data available for model and non-model organisms to estimate species trees, which is revolutionizing the field of molecular phylogenetics (Edwards, 2009).

Recently, a phylogenomic dataset of the class Insecta was generated to estimate the phylogeny and divergence times for the major orders of this lineage (Misof et al., 2014). However, gene tree heterogeneity was not considered when reconstructing the phylogenetic relationships. Compared to other biological groups, the Insecta poses a major challenge to MSC-based inference. This is mainly assigned to the large effective population sizes (Romiguier et al., 2014) associated with short speciation intervals, which leads to high degrees of ILS (Maddison and Knowles, 2006; Pamilo and Nei, 1988). Further, this class still has several controversial relationships, e.g., the Strepsiptera position within Neuropteroidea (Boussau et al., 2014; Huelsenbeck and Biology, 2007; Whiting et al., 1997), the position of Paraneoptera (Kjer, 2004; Kjer et al., 2006), the interordinal relationships within Polyneoptera (Whitfield and Kjer, 2008), and the sister-group of Insecta (Regier et al., 2010; Trautwein et al., 2012).

Once the MSC approach was already successfully applied to resolve deep phylogenies (Song et al., 2012; Xi et al., 2014), we were prompted to test how this methodology would impact the interordinal relationships of Insecta. Since large ancestral population sizes coupled with consecutive speciation events may cause high discordance between gene and species trees, the species tree inferred by MSC should better reflect lineage history. Therefore, we expect that interordinal branching, which are deeper located in the phylogeny, would present a distinct pattern when compared with the supermatrix based approach. To test this, we estimated the phylogenetic relationships of Hexapoda using the MSC approach and a genomic dataset, focusing on the interordinal relationships within this subphylum. We found three main differences compared with the supermatrix approach: the non-monophyly of Hexapoda and the interordinal relationships within Polyneoptera and Neuropteroidea, including the non-monophyly of Neuroptera, which indicates that more studies are urged to better elucidate the evolutionary patterns of this lineage.

## Material and methods

We downloaded all amino acid alignments (1,478 genes) used to reconstruct the phylogenetic relationships of Hexapoda, as well as the concatenated supermatrix C from the dataset of (Misof et al., 2014), available from the Dryad repository (http://dx.doi.org/10.5061/dryad.3c0f1). Then we used RAxML v. 8.2.6 (Stamatakis, 2014) to estimate the ML (maximum likelihood) gene trees with the LG+G(4) model of amino acid substitution (Le and Gascuel, 2008; Yang, 1993). For each of the 1,478 genes, 100 bootstrap replicates were inferred using CIPRES REST API (Miller et al., 2015). (S1 file).

After gene tree estimation, ASTRAL-II v. 4.10.2 (Sayyari and Mirarab, 2016) was used to independently reconstruct two species trees. Firstly, the complete dataset with 1,478 gene trees was used to estimate the Hexapoda phylogeny. To investigate the effects of gene sampling, the species tree was estimated using 377 gene trees inferred from alignments containing more than 500 amino acids. This dataset with longer genes was composed to minimize the effects of sampling errors in gene tree topology estimation, which arguably lead to a higher probability of inferring the correct species tree (Song et al., 2012). In ASTRAL, 100 bootstrap replicates were estimated to obtain branch supports.

To evaluate the robustness of bootstrap values of the species tree, we used the ape package v. 3.5 (Paradis et al., 2004) of R language v. 3.2.0 (R Core Team, 2016) to draw a correlation between the bootstrap support value and the number of genes and sites in the alignments. Thus, we investigated whether nodes with low bootstrap support were the ones that included species with lower number of genes/sites. Additionally, Robinson-Foulds distances were computed comparing the inferred gene trees to Mifof’s phylogeny and to both species trees that we inferred in ASTRAL (with all genes and with genes that were longer than 500 amino acids).

## Results and Discussion

Our inferred species tree from the full dataset failed to support the monophyly of Hexapoda, as recovered by Misof et al. (Misof et al., 2014), due the placement of a crustacean species, the remipede *Speleonectes tulumensis*, as the sister-group of Insecta (Figure 1). This result was also recovered in the dataset containing gene alignments longer than 500 bp (Figure S1). The only interordinal difference within Hexapoda between both inferred species trees was that the Megaloptera was recovered as monophyletic only when all genes were used. Also, in both datasets, MSC failed to infer the monophyly of Neuroptera. *Conwentzia psociformis* was placed as sister group to the remaining Neuropterida (Raphidioptera + Megaloptera + Neuroptera) in the complete dataset and as sister-group to the Neuropterida (with exception of *Sialis*) in the dataset with longer alignments.

**Figure 1:**
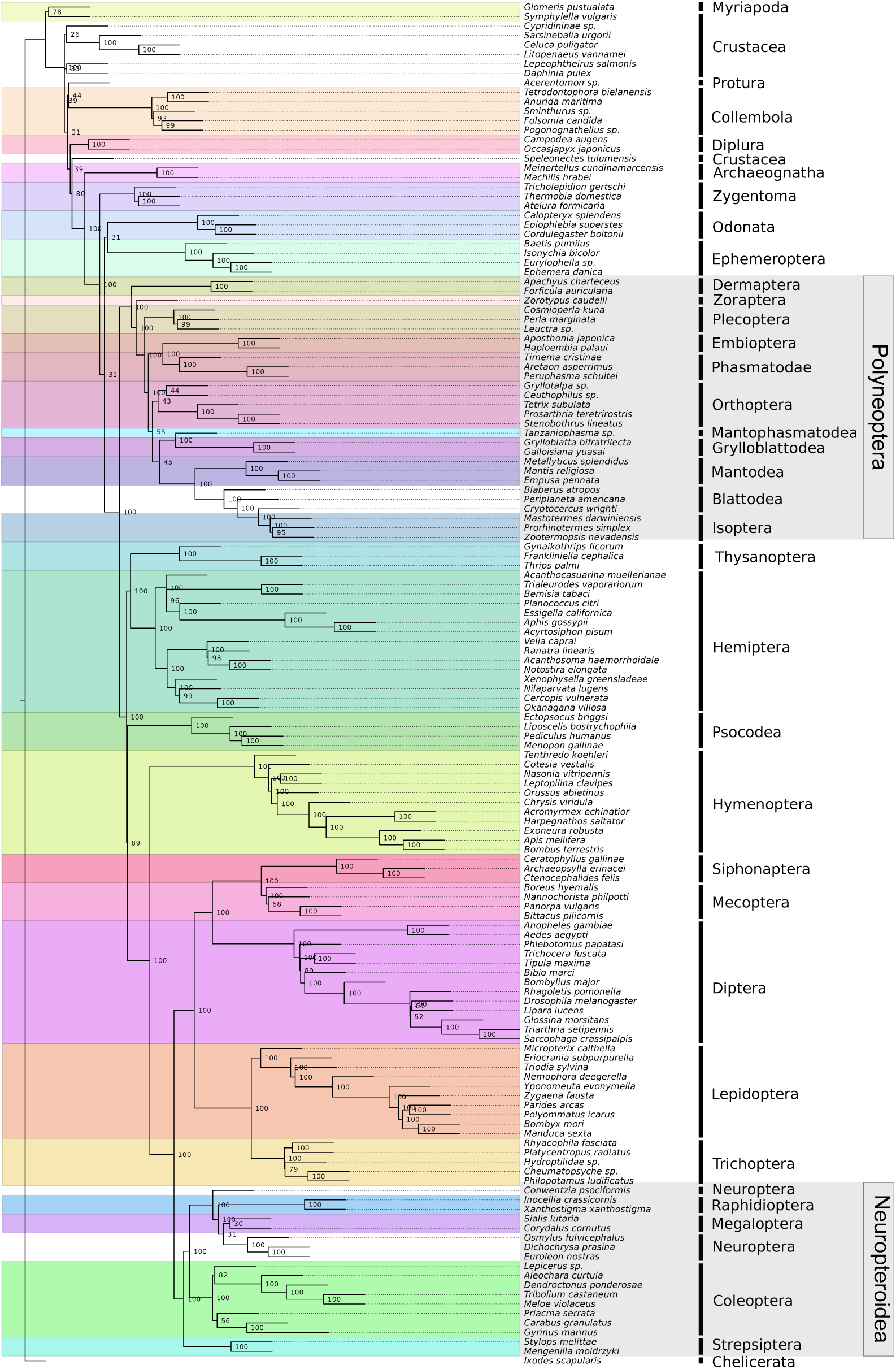
Insect species tree inferred from the full dataset. Support values on nodes were obtained from a hundred bootstrap replicates. Distinct colors indicate ordinal clades. Gray boxes indicate the different interordinal relationships when compared to Misof et al. (2014)’s results.

In Insecta, all orders except Megaloptera, Neuroptera and Blattodea were inferred as monophyletic by both data sets. Moreover, several major supra orginal groups were also recovered as monophyletic, e.g., Holometabola, Condylognatha, Polyneoptera and Palaeoptera. The phylogenetic relationships within Hexopoda orders were stable between data sets, with the exception of Coleoptera. In the complete dataset, *Priacma* was inferred as sister-clade of a group formed by *Carabus* and *Gyrinus*; this three-species clade was the sister-group of the remaining Coleoptera. When larger alignments were analyzed, *Priacma* was the sister-group of the remaining Coleoptera.

### Non-monophyly of Hexapoda

As (Misof et al., 2014), Our phylogenetic tree also recovered a clade formed by Protura and Collembola, which is the sister-group of all the remaining Hexapoda (Diplura, Insecta) in this same work. Our species tree did not recover Misof et al.’s hexapod arrangement, ((Protura, Collembola), (Diplura, Insecta)), because the Remipedia crustacean *Speleonectes tulumensis* was inferred as the sister-group of Insecta, whereas Diplura was the sister-group of *S*. *tulumensis* + Insecta. Von Reumont et al. (Von Reumont et al., 2012), using phylogenomic data, also found a clade composed of Protura and Collembola, in their analysis, however, Diplura was the sister-group of Protura and Collembola, whereas Insecta was the sister-group of these three clades.

The majority of recent molecular phylogenetic surveys of Hexapoda proposed the arrangement (((Diplura, Protura), Collembola), Insecta) (Kjer et al., 2006; Meusemann et al., 2010; Rainford et al., 2016; Trautwein et al., 2012), although only Meusemann et al. (Meusemann et al., 2010) used a phylogenomic dataset to infer the phylogeny. An alternative higher order evolutionary relationship of hexapods was propoded by Sasaki et al. (Sasaki et al., 2013), who estimated the topology (((Collembola, Diplura), Insecta), Protura). When compared to Misof et al.’s (Misof et al., 2014) tree, our species tree presented few interordinal differences within Insecta, mainly associated with the Neuropteroidea and Polyneoptera supergroups.

### Weak support for Neuropteroidea

In the consensus phylogeny of Neuropteroidea, this lineage is composed by Neuropterida – ((Megaloptera, Neuroptera), Raphidioptera), which is sister to the (Coleoptera, Strepsiptera) clade (Mckenna et al., 2015; Misof et al., 2014; Peters et al., 2014; Wiegmann et al., 2009). In our analysis, relationships within Neuropterida differed. While recent studies supported either the previous topology (Cameron et al., 2009; Mckenna et al., 2015; Misof et al., 2014; Peters et al., 2014; Wang et al., 2012; Wiegmann et al., 2009; Yan et al., 2014; Zhao et al., 2013), or the remaining two topological alternatives (Aspöck et al., 2012; Ishiwata et al., 2011; McKenna and Farrell, 2010), our results supported the non-monophyly of Neuroptera for the first time using molecular data, placing *Conwentzia psociformis* as sister-group to the remaining Neuropterida.

It is worth mentioning that in previous studies using morphology or molecules the monophyly of Neuroptera was undisputed (Aspöck et al., 2012). Most studies, however, used only one to three Neuroptera species, and a few genes that were concatenated into a single supermatrix for phylogeny inference. Thus, poor taxonomic sampling – the highest number of Neuroptera species included in a single analysis was 10 (Yan et al., 2014), associated with low number of sampled genes and the handling of data likely led to differences regarding Neuroptera relationships. The bootstrap support value for the Neuropterida clade excluding *Conwentzia psociformis* was very low (31). Such a low support indicates that additional genomic data from other species of these groups are needed to better resolve this node (Lambert et al., 2015).

The second issue within Neuropteroidea was the position of the Strepsiptera order, which has been debated for a long time (Beutel and Pohl, 2006; Whiting et al., 1997). The consensus based on recent works put this order together with Coleoptera (Boussau et al., 2014; Cameron et al., 2009; Mckenna et al., 2015; Misof et al., 2014; Peters et al., 2014; Wang et al., 2012; Wiegmann et al., 2009; Yan et al., 2014; Zhao et al., 2013), however our results do not corroborate to this general view, since Strepsiptera was placed as sister-group to all others Neuropteroidea with high support (100). This way, our findings indicate that Strepsiptera problem is still open, bringing up more discussion on this relationship.

### Polyneoptera interordinal relationships

Within the Polyneoptera superorder, differences between Misof et al.’s (Misof et al., 2014) and the species trees estimated in the present work were more relevant. In our species tree, Orthoptera was the sister-group of a clade formed by Mantophasmatodea, Grylloblattodea, Mantodea, Blattodea and Isoptera with bootstrap support value of 55. This six-order clade clustered with a group composed of Empioptera and Phasmatodea (100). In Misof et al.’s tree, the Orthoptera was placed in basal position within this lineage, with Mantophasmatodea and Grylloblattodea clustering with Embioptera and Phasmatodea. This four-order clade was clustered with Dictyoptera (Mantodea, Blattodea and Isoptera), which was also recovered in our species tree. The remaining differences within the Polyneoptera was the placement of Zoraptera and Demaptera. In Misof et al.’s tree these two orders were clustered together and they were the sister-group of the rest of Polyneoptera, while in our species tree Demaptera was the sister to the remaining Polyneoptera.

A general comparison with previous works is complicated by the fact that the number of orders used varied significantly among studies, making a one-to-one evaluation of the phylogenetic placement of Polyneoptera orders difficult. Although previous studies employed a varied combination of taxon and genes, they estimated some of the clades found here; for instance, the Dictyoptera (Ishiwata et al., 2011; Ma et al., 2014; Misof et al., 2014; Sasaki et al., 2013; Tomita et al., 2012; Wan et al., 2012; Wu et al., 2014), the Embioptera + Phasmatodea (Ishiwata et al., 2011; Misof et al., 2014; Sasaki et al., 2013; Wu et al., 2014) and the Mantophasmatodea + Grylloblatodea (Ma et al., 2014; Misof et al., 2014; Sasaki et al., 2013).

### Support values of controversial relationships

It is worth noting that most nodes with poor support lied within Neuropteroidea and Polyneoptera groups. As shown by Lambert et al. (Lambert et al., 2015), topologies estimated by MSC methods and concatenation will tend to disagree when short branches and low support values are present in the concatenated tree. Differences between our species tree and Misof et al.’s concatenated tree possibly recapitulated this pattern. In Misof et al.’s concatenated tree, there were 7 support values below 50, with 3 located at interordinal ancestral nodes. These three nodes resulted in interordinal discordances between Misof et al.’s (Misof et al., 2014) and our MSC trees, namely, (i) the support of (Protura, Collembola) as sister-group of (Diplura, Insecta) in Misof et al’.s tree was 14, whereas our support for *Speleonectes* as sister-group of Insecta was 80; (ii) the support for Dictyoptera as sister-group to the ((Mantophasmatodea, Grylloblattodea), (Embioptera, Phasmatodea)) clade in Misof et al.’s (Misof et al., 2014) tree was 34 and we inferred Dictyoptera as sister-group of (Mantophasmatodea, Grylloblattodea) with bootstrap support value of 45; (iii) the bootstrap support of for the relationship of Neuroptera and Megaloptera as sister-groups was 15 in Misof et al.’s tree, whereas we recovered Neuroptera as non-monophyletic clade with bootstrap value of 31.

Additionally, we investigated whether a correlation existed between the number of genes (or sites) (S1 Table) and the bootstrap support values of the species tree, but no significant correlation was found for either the number of genes or sites. Because no correlation was found between the number of sites and the bootstrap support, it is unlikely that sequencing more genes alone will solve the remaining controversies on insect phylogeny. Instead, it may be possible that increasing the number of sampled species for the orders near poorly-supported clades would improve topological support. The addition of taxa to clarify the discordances was already recommended in other works as well.

Given the discordances between our MSC tree and Misof at al.’s tree, we were prompted to investigate which of the two topologies lied closer to the forest of inferred gene trees. To this end, we calculated the distribution of topological distances between Misof et al.’s and the MSC tree and the gene tree forest. The distribution of RF distances failed to indicate any departure between both topologies tested, implying that distance from the forest of gene trees does not favor any topological hypothesis (Figure 2).

**Figure 2:**
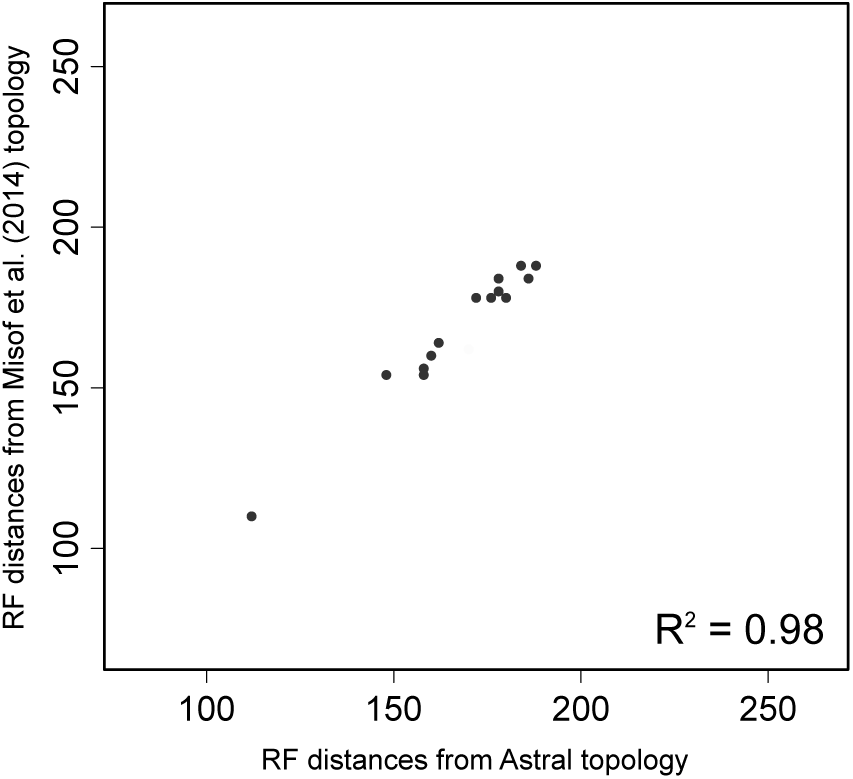
Robinson-Foulds distance values computed between each gene tree and Misof et al. (2014) topology and Astral topology. The dashed blue line indicates the one to one relationship.

### Species tree

Despite some criticism on the MSC method (Springer and Gatesy, 2016), it has clear advantages over concatenated methods, *e. g.* better performance in the presence of high degrees of ILS and short branches and the incorporation of gene tree heterogeneity (Edwards et al., 2016; Liu et al., 2015). Then, using a huge data (144 species and 595,033 amino acid sites) we could infer a deep-level phylogenetic tree under the MSC approach. The comparison of the recovered species tree with phylogenetic hypotheses based solely on concatenated methods showed three main disagreements: the non-monophyly of Hexapoda, which also revealed an incongruence at the root of Insecta, and the interordinal relationships within Polyneoptera and Neuroptera.

Even with the challenges caused by the great size of the effective population size associated with short branches in Insecta, our MSC analysis was able to recover several orders and relationships according to the literature of Insecta systematics. Further, the results and discussion presented here clearly show the most incongruent nodes are in the Insecta tree of life, for which more studies are necessary to achieve a better historical hypothesis for the group.

## Acknowledgments

This work is part of the Doctoral thesis of LAF funded by Brazilian Ministry of Education (CAPES) and the Brazilian Research Council (CNPq).

---

**Figure S1.**
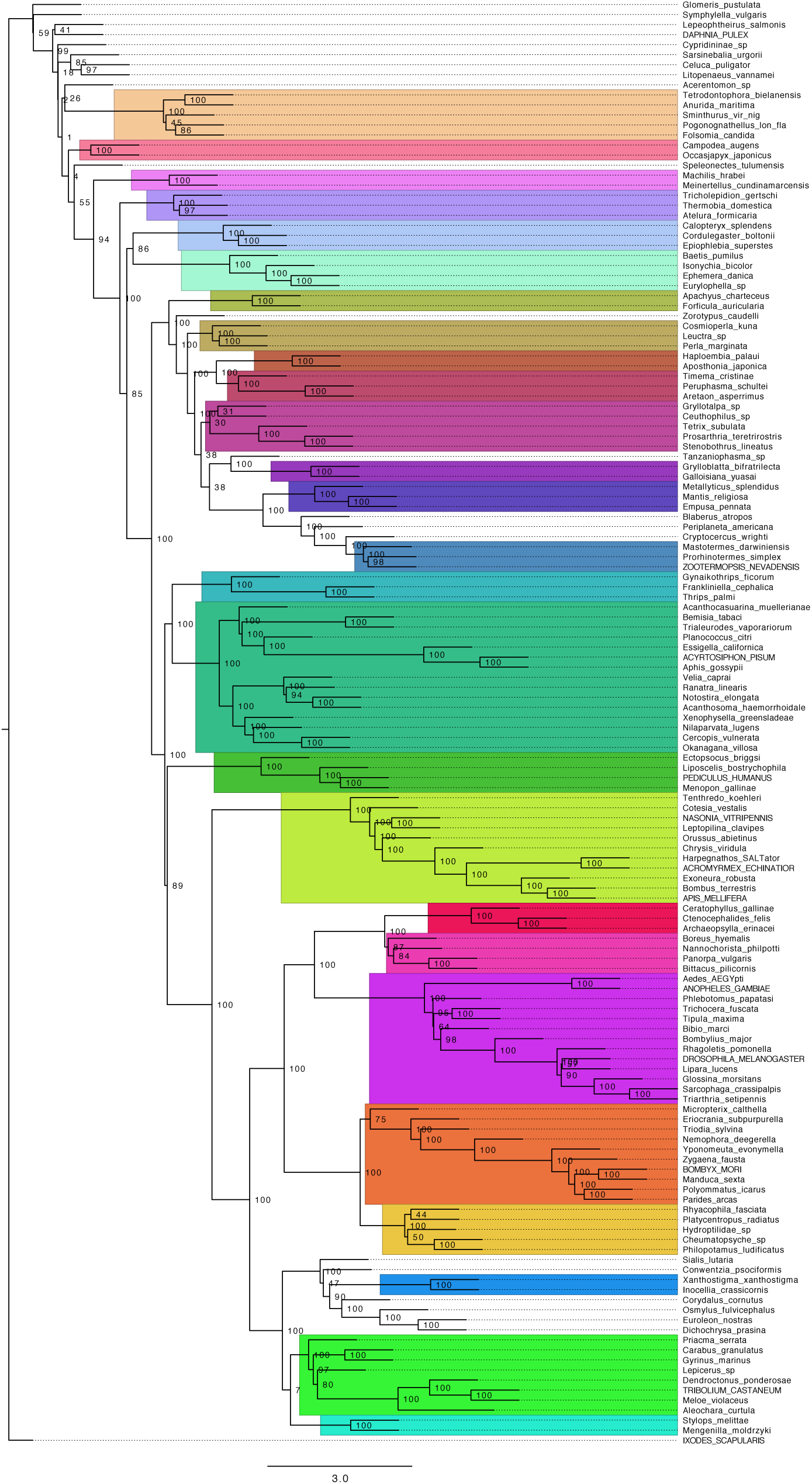
Phylogenetic tree inferred using 377 genes with more than 500 amino acids

